# The impact of heat on human physical work capacity; part III: the impact of solar radiation varies with air temperature, humidity, and clothing coverage

**DOI:** 10.1101/2021.07.01.449782

**Authors:** Josh Foster, James W Smallcombe, Simon Hodder, Ollie Jay, Andreas D Flouris, Lars Nybo, George Havenith

**Author notes:** Correspondence: Professor George Havenith, Environmental Ergonomics Research Centre, Loughborough University, Loughborough, LE11 3TU, Leicestershire, UK, E.

## Abstract

It is well-known that heat impacts human labour/physical work capacity (PWC), but systematic evaluations of solar radiation (SOLAR) effects and the interaction with air temperature and humidity levels and clothing are lacking, as most lab-studies are conducted in semi-nude subjects without radiation or only in a single climatic condition. Due to the high relevance of SOLAR in various occupations, this study quantified how SOLAR interacts with clothing and other primary environmental factors (air temperature/humidity) of importance to determine PWC in the heat. The data allowed the development of a SOLAR correction factor for predicting PWC in major outdoor industries. Fourteen young adult males (7 wearing a standardised work coverall (0.9 Clo), 7 with shorts and trainers (0.3 Clo) walked for 1-hour at a fixed heart rate of 130 b·min^−1^, in seven combinations of air temperature (25 to 45°C) and relative humidity (20 or 80%), with and without SOLAR (800 W/m^2^ from solar lamps). Cumulative energy expenditure in the heat, relative to the work achieved in a cool reference condition, was used to determine PWC%. Skin temperature was the primary determinant of PWC in the heat. In dry climates with exposed skin (0.3 Clo), SOLAR caused PWC to decrease exponentially with rising air temperature, whereas work coveralls (0.9 Clo) negated this effect. In humid conditions, the SOLAR-induced reduction in PWC was consistent and linear across all levels of air temperature, and clothing conditions. WBGT and UTCI based prediction equations of PWC represented SOLAR correctly. For heat indices not intrinsically accounting for SOLAR, correction factors are provided enabling forecasting of heat effects on work productivity.

## Introduction

Environmental heat exposure has a negative impact on human health and physical working capacity (PWC) (Flouris et al., 2018; Foster et al., 2021b, 2021a; Ioannou et al., 2021), incurring significant economic damage through its impact on workplace productivity (Hsiang et al., 2017; Hübler et al., 2008; Zander et al., 2015). Understanding the full effect of heat on PWC is required for economic cost and general impact analysis associated with climate change and hot weather events in general (Hsiang et al., 2017). While models of PWC based on various climate indices have recently been developed (Dunne et al., 2013; Foster et al., 2021b; Kjellstrom et al., 2018), at present, none account for the effect of solar or general thermal radiation. Given that many occupational tasks involve outdoor exposure, not accounting for thermal radiation is presently a significant limitation.

In 338 trials with work paced based on heart rate (limit of 130 b·min^−1^, [moderate to heavy work]) our group recently developed empirical models for PWC based on a suite of heat stress indices (Foster et al., 2021b). However, those trials were conducted without added solar radiation (SOLAR). Hence, to use the equations for both conditions with and without SOLAR, correction factors may be needed, especially for heat stress indices that do not intrinsically account for this parameter. For example, although wet bulb temperature (*T*_wb_), Humidex, and Heat Index strongly predict PWC in shaded environments (Foster et al., 2021b), they are calculated from *T*_a_ and Rh alone, and therefore cannot accommodate conditions in which there is additional SOLAR. Moreover, while wet-bulb globe temperature (WBGT) and universal thermal climate index (UTCI) account for radiation in their calculation (Havenith and Fiala, 2015), their correct sensitivity to radiation must also be validated empirically. For WBGT and UTCI, one equation linking PWC to the thermal climate would be expected since they account for radiation, whereas correction factors may be required for *T*_wb_, Humidex, and Heat Index.

Thermal radiation is electromagnetic radiation emitted from hot surfaces, with the precise waveband characteristics dependent on the surface temperature of the emitting material (Miller, 2012). When electromagnetic radiation is absorbed and retained by the human body, it increases body heat content which elevates thermal and cardiovascular strain (Bröde et al., 2008, 2010a). Studies involving outdoor activity in the sun show marked elevations in thermal strain compared with shaded conditions, for the same air temperature (*T*_a_) (Adolph, 1947; Gonzalez et al., 2012; Hardy and Stoll, 1954; Nielsen et al., 1988; Otani et al., 2017, 2019). The detrimental effects of radiation on thermal strain (Gagge and Hardy, 1967; Nielsen, 1990; Stolwijk and Hardy, 1965), maximal exercise performance (Otani et al., 2016), cognitive function (Piil et al., 2020), and thermal comfort (Hodder and Parsons, 2007) have also been investigated with SOLAR lamps, a source of artificial, non-ionising radiation.

Given the interaction between SOLAR and the level of human heat strain, exposure to sunlight during physical work is likely to reduce PWC. However, the extent to which SOLAR influences PWC across wide variations of *T*_a_ and relative humidity (Rh) is unknown. For example, the impact of SOLAR on PWC may depend on the pre-existing climate, or, presuming a fixed solar load, may be consistent regardless of the environment. Moreover, it is unknown to what extent protective clothing alters this response. While clothing colour and fabric can have a significant impact on overall solar heat gain (Bröde et al., 2008, 2010b; Nielsen, 1990), direct comparisons between exposed and clothed skin are unavailable. The above knowledge is essential for accurate PWC forecasting, and to determine what combinations of *T*_a_ and Rh should be targeted with shading solutions and clothing modifications.

The primary aim of this study was to quantify the impact of SOLAR on PWC, based on clothing coverage (low or high), across a wide range of temperature and relative humidity combinations (*T*_a_:25-45C Rh: 20-80%). The secondary aim was to generate correction factors so PWC can be modelled outdoors using heat stress indices that do not ordinarily account for SOLAR in their calculation (namely *T*_wb_, Humidex, and Heat Index). The final aim was to test the sensitivity of WBGT and UTCI to added solar heat loads, confirming whether our earlier published functions (Foster et al., 2021b) can be used for indoor and outdoor work settings.

## Methodology

### Overview

Participants were randomly allocated into either a low or high clothing coverage group (*between participant*), and subsequent *within participant* comparisons (i.e., SOLAR vs SHADE) took place in different *T*_a_ and Rh combinations. We adopted a physical work protocol in a climate chamber which simulates the self-pacing behaviours adopted in the field. We refer the reader to our companion paper (Foster et al., 2021b) for a thorough rationale and description of the protocol. A study schematic is shown in Figure 1.

**Figure 1.**
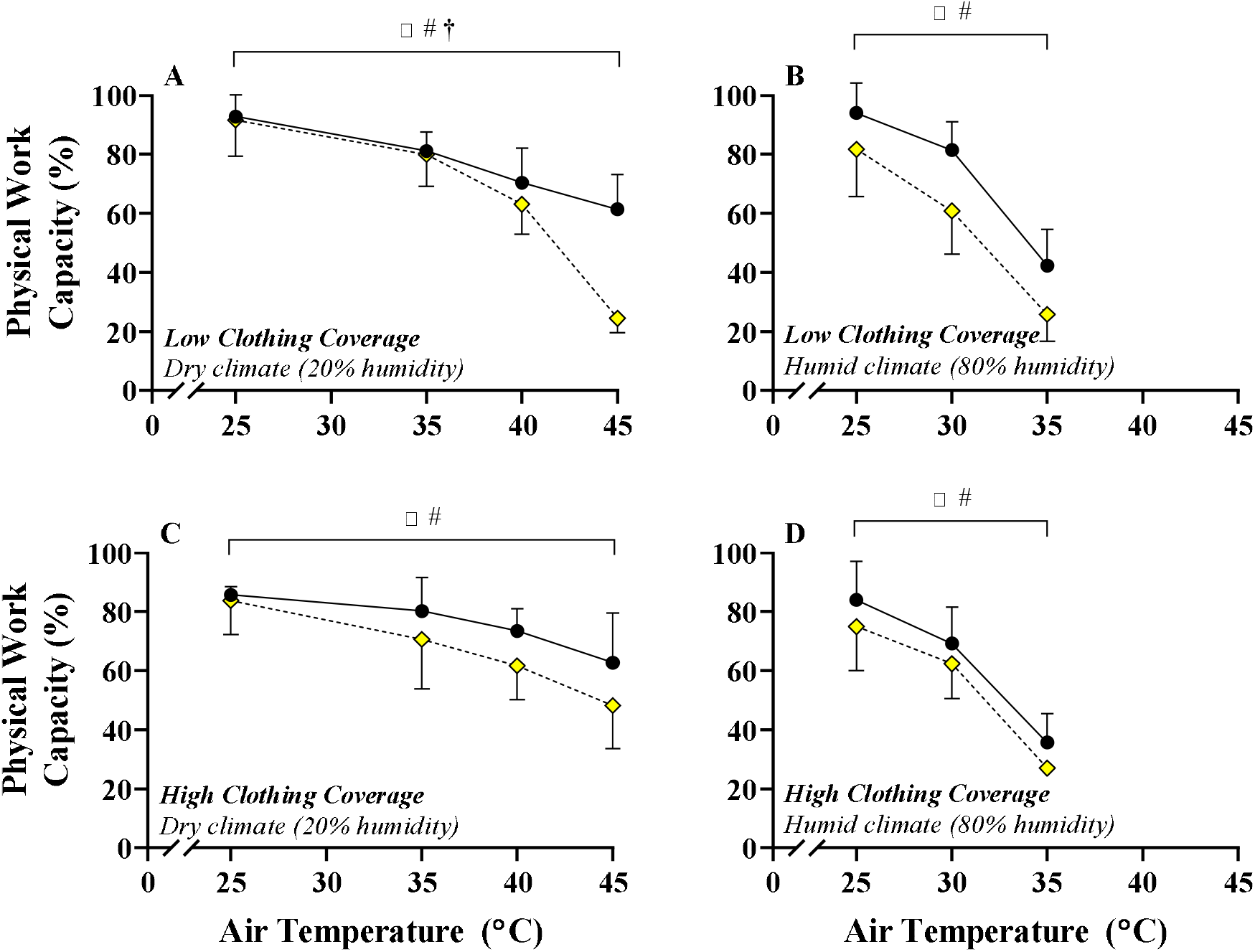
Change in absolute physical work capacity in SHADE (black circles) with SOLAR (yellow diamonds) in low (A and B) or high (C and D) clothing coverage conditions, in dry (A and C) and humid (B and D) climates. ⍰**#**denote main effects for condition and air temperature. **†**denotes an interaction effect between condition and air temperature.

### Ethical approval

This study was approved by the Loughborough University Ethics Committee and was performed in-line with the Declaration of Helsinki. Participants were provided with an information sheet that detailed the risks and requirements of the experiment before providing written informed consent. Participants conducted a health screening questionnaire prior to the start of the experiments.

### Location and timeline

The data collection took place in custom made environmental chambers (TISS performance chambers, UK) located within the Environmental Ergonomics Research Centre, Loughborough University. Data collection ran from September 2018 to May 2019.

### Participants

A total of thirteen participants took part in the study. The total number of trials performed varied between participants, ranging from 4-14 (median = 10). Table 1 displays participant characteristics for each experimental group. To reduce any impact of heat acclimation throughout the trial period, the number of experiments were capped at three per week, but seldom exceeded two per week. The fixed heart rate approach also limits any sustained rises in body temperature that are required on a daily basis to elicit a heat acclimated phenotype (Foster et al., 2021b; Fox et al., 1963). Moreover, participants were not permitted to take part in the study if they recently visited a hot climate.

**Table 1.** Participant characteristics. Data are presented as means ± standard deviation. The data range are presented in parentheses.

| Variable | Low Clothing Coverage<br>(n = 7) |  | High Clothing Coverage (n = 7) |  |
| --- | --- | --- | --- | --- |
| Age (years) | 25 ± 3 | (20 - 29) | 23 ± 3 | (20 - 28) |
| Height (cm) | 177 ± 5 | (171 - 185) | 178 ± 6 | (172 - 192) |
| Mass (kg) | 74 ± 10 | (59 - 92) | 74 ± 11 | (59 - 94) |
| Body Surface Area (m<sup>2</sup>) | 1.9 ± 0.1 | (1.7 - 2.2) | 1.9 ± 0.1 | (1.7 - 2.1) |
| Body Mass Index (kg·m<sup>-2</sup>) | 23 ± 2 | (20 - 28) | 23 ± 3 | (20 - 29) |
| Body fat (%) | 18 ± 6 | (10 - 27) | 14 ± 4 | (9 - 21) |
| $\dot{V}\text{O}_{2\text{max}}$ (L·min<sup>-1</sup>) | 3.8 ± 0.8 | (2.6 - 5.9) | 4.2 ± 0.0 | (2.7 - 5.3) |
| $\dot{V}\text{O}_{2\text{max}}$ (mL·kg<sup>-1</sup>·min<sup>-1</sup>) | 52 ± 9 | (40 - 65) | 56 ± 5 | (44 - 64) |

$\dot{V}\text{O}_{2\text{max}}$ , maximal oxygen consumption; BMI, body mass index; BSA, body surface area; L, litres; mL, millilitres;

### Experimental Design

Thirteen young adult male participants were allocated to either a low or high clothing coverage group. One participant completed trials in both clothing conditions, totalling n=7 in each. Participants first completed a graded exercise test on a treadmill to determine maximal oxygen consumption (V◻O_2max_). Participants then completed 1 hour of walking exercise at a fixed heart rate of 130 b·min^−1^, first in a reference cool condition, and thereafter in up to seven different combinations of *T*_a_ and Rh. In terms of the *T*_a_ and Rh combination, trials were completed in a randomised order, but a solar vs. no solar comparison trial was always completed sequentially (i.e., one followed the other, though always on separate days). A total of 66 and 62 hot trials were completed in low and high clothing coverage, respectively. In the supplementary file, Table S1 displays the number of trials performed by each participant according to the air temperature and relative humidity combination. A study schematic is shown in the supplementary file (Figure S1).

### Experimental controls

Participants completed experimental sessions at the same time of day to minimise the effect of circadian rhythm on outcome variables (Waterhouse et al., 2004). However, it is worth noting that, apart from changes in absolute core temperature, physiological effector responses are unaffected by time of day (Ravanelli and Jay, 2020). Participants presented to the lab in a hydrated state (confirmed by urinary analysis), and refrained from caffeine 12 hours prior to each trial. Finally, participants were asked to refrain alcohol and vigorous exercise 24 hours before each trial.

### Preliminary trial (visit 1)

The preliminary visit involved an anthropometric assessment and a submaximal (walking) test of maximal oxygen consumption (V◻O_2max_) performed on a treadmill. Body composition was assessed using a Tanita scale (MC-780MA, TANITA Corporation, Japan) while participants were dressed in underwear only. At a fixed walking speed of 4.5 km/hour, the submaximal test followed a ramp protocol in which the gradient increased by 5% every 3 minutes until a steady state heart rate of 85% age predicted maximum was attained. Combined with indirect calorimetry to continuously assess V◻O_2_ uptake, V◻O_2max_ was predicted by extrapolating V◻O_2_ to age predicted maximum heart rate. The protocol is set out in more detail in our companion paper (Foster et al., 2021b).

### Experimental protocol

Upon arrival, participants inserted a rectal thermistor (VIAMED, Yorkshire, UK) to a depth of 10 cm past the anal sphincter to monitor internal (rectal) temperature, which allowed continuous monitoring of core temperature throughout each trial. Participants subsequently voided their bladder and provided a urine sample which was used for the assessment of urine specific gravity. If urine specific gravity exceeded 1.020, participants drank 500 ml of water and provided another sample after 20 minutes (Armstrong et al., 1994). Skin thermistors (Grant Instruments Ltd, Corby, UK) were then placed onto each participant at 6 sites, allowing continuous measurement of skin temperature (*T*_skin_). Thermistors were attached to the following sites with a breathable, hypafix tape (BSN medical, D-22771, Hamburg, Germany); the upper back, lower back, chest, arm (triceps), thigh (quadriceps), and calf. The mean *T*_skin_ was then calculated by adapting the Ramanathan equation (1964). The original equation is equal to:

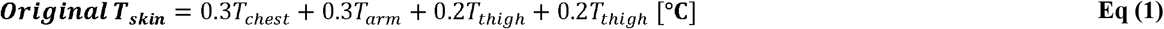

To integrate additional skin temperature sites in the radiated area at the upper and lower back, *T*_chest_ was replaced by *T*_torso_, providing equal weight to the front and back temperatures of the torso, as below:

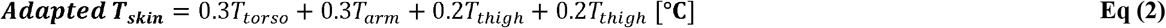

Where *T*_torso_ was calculated by:

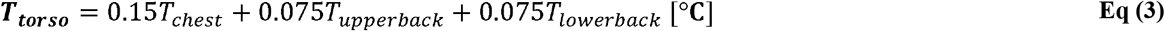

The value for mean *T*_skin_ was reported as the average score of “adapted *T*_skin_” during a 1-hour trial.

### Physical work simulation

The treadmill was programmed to control workload to achieve the desired heart rate of 130 b·min^−1^. The treadmill speed and grade were never manually controlled by the researchers or participants. The treadmill incline did not change until the speed reached its maximum of 6 km·h^−1^. Thereafter, the speed did not change unless the incline fell back to zero. Each test was set to last a maximum 1 hour, but in more extreme heat the speed often fell to zero before this time. If speed fell to zero, (i.e., resting heart rate is ≥ 130 b·min^−1^) the participants exited the chamber and no more exercise took place.

### Calculation of percentage physical work capacity

A predictive equation based on treadmill speed and grade (Ludlow and Weyand, 2017) was used to calculate total cumulative energy generated in the present study. The original equation was expanded to convert energy generated in V◻O_2_ to total kilojoules. This equation, and its validation based on 365 expired air samples are available in our companion paper (Foster et al., 2021b).

### Clothing

Participants were separated into two clothing groups before undertaking trials with and without SOLAR radiation. The two conditions were chosen to provide minimal/low or high clothing coverage of the skin. In the low clothing trials, subjects wore underwear, standardized shorts, socks, and trainers. In the high clothing trials, subjects wore the same as the low clothing trial, with the addition of a standardized cotton t-shirt, and a standardized full body protective coverall (65% polyester, 35% cotton). The intrinsic clothing insulation of the low and high coverage ensembles were estimated as 0.04 and 0.133 m^−2^·K·W^−1^ (0.26 and 0.86 Clo), respectively, based on the International Standard (ISO9920, 2009). Using the equation provided in the standard, the evaporative resistance was estimated at 0.007 and 0.024 m^−2^·kPa·W^−1^ for the low and high clothing conditions, respectively.

### Environmental logging

A Quest-temp model 34 meter was used to record wet-bulb globe temperature (WBGT) at 1-minute intervals. The approach taken to measure WBGT is described below. A Testo model 435-2 with hot-wire probe was used to record *T*_a_ (shaded), Rh, and air velocity and logged at 1-minute intervals.

### Calculation of Heat Stress Indices

#### Wet bulb globe temperature (WBGT)

For all shaded trials, the average value for dry bulb, wet bulb, and globe temperature were used to assess WBGT (WBGT= 0.7 *T*_wb_+0.2*T*_globe_+0.1*T*_a_) for a given work bout. For the SOLAR trials, live WBGT measurement was not practical since the WBGT meter needed to be placed in the same location as the exercising human participant, and at different heights (radiation intensity varied with height). Hence, the value for natural wet-bulb and globe were measured at four heights on a separate occasion while exposed to SOLAR. The dry bulb was measured in each SOLAR trial, shielded from radiation. Therefore, the same value for natural wet bulb and globe temperature was used for each participant for each *T*_a_ and Rh combination with SOLAR.

#### Psychrometric wet bulb temperature (*T*_wb_)

Psychrometric wet bulb temperature was calculated based on Bernard & Pourmoghani (1999):

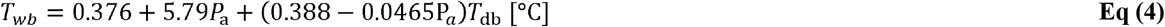

Where *P*_a_ is the ambient water vapour pressure measured in kPa, and *T*_db_ is the dry bulb temperature (air temperature). P_a_ was calculated using the following equation (Parsons, 2010):

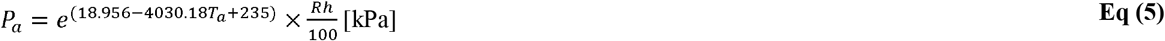

Where *T*_a_ is ambient temperature in °C, and Rh is relative humidity (0-100).

#### Universal Thermal Climate Index (UTCI)

UTCI was calculated for each condition in excel (www.climatechip.org/excel-wbgt-calculator), which uses a 6 parameter polynomial model (Bröde et al., 2012). *T*_a_, Rh, globe temperature and air velocity were used for the calculation.

#### Humidex

The Humidex was computed as below (Masterton and Richardson, 1979; Rana et al., 2013):

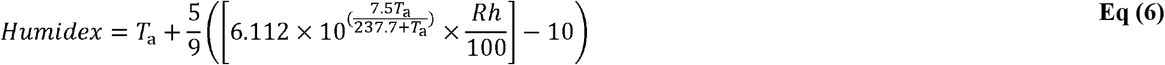

Where *T*_a_ is air temperature in degrees Celsius; *Rh* is relative humidity in % (0-100).

#### Heat Index

The Heat Index was computed as below (Rothfusz, 1990):

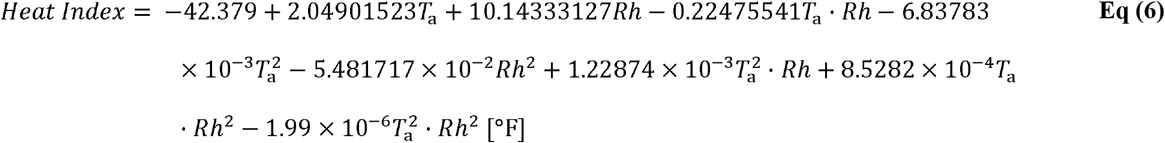

Where *T*_a_ is in degrees Fahrenheit and Rh is 0-100. The Heat Index in Fahrenheit (HI_F_) was converted to degrees Celsius by:

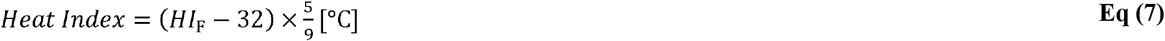

### Solar simulation lamps

SOLAR was produced artificially using compact source iodide lamps (CSI, Thorn Lighting, Durham, UK; (Beeson, 1978). The lamps filter out ionising radiation to negligible values, and thereafter produce a similar spectral content to that of sunlight. An array of three vertically aligned 1,000 W metal halide lamps were placed 2.3 m posterior to the participant. In the SOLAR trials, the lamps were switched on at least 1-hour before activity. The average intensity of radiation across all exposed regions for all trials was 807 ± 24 W/m^−2^, as measured immediately before and after each experiment (Pyranometer, Kipp and sons, The Netherlands). This level of SOLAR is typical for what is observed under a clear sky during the hottest part of the day (Ioannou et al., 2017; Monteith and Unsworth, 1990). The projected area (A_p_) was estimated at 24.2% total body surface area, based on a SOLAR altitude and azimuth of 0° (Underwood and Ward, 1966). The average body surface area was 1.90 m^−2^ (range 1.69 to 2.21 m^−2^), resulting in a total radiant heat load of 382 Watts (range 339 to 441 Watts).

### Statistical Analysis

Statistical analyses were conducted using IBM SPSS version 27 and GraphPad Prism version 8. In SPSS, a linear mixed model with fixed [condition (SOLAR or SHADE), temperature (three or four levels depending on humidity), and clothing (low or high coverage)] and random (subject ID) effects was used to compare physical work capacity (PWC%) responses between SHADE and SOLAR trials. Four levels of temperature were assessed in dry climates (25, 35, 40, and 45°C), and three were assessed in humid climates (25, 30, and 35°C). Data are reported as mean difference ± standard error, 95% confidence intervals of the difference, and effect size. Effect size was calculated as:

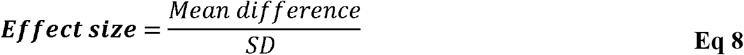

where mean difference is the group average difference between PWC in SHADE vs SOLAR, and SD is the standard deviation of the differences. The threshold values of 0.2, 0.5, and 0.8 were used to indicate a small, moderate, and large effect, respectively (Cohen, 1988).

In accordance with our companion paper (Foster et al., 2021b) we used a sigmoidal expression to determine the impact of heat on PWC%. The model takes the form:

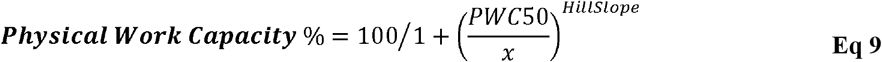

Where *x* represents the predictor studied (e.g. WBGT), *PWC50* is the value of *x* that elicits 50% PWC, and *HillSlope* defines the steepness of the curve. The *HillSlope and PWC50* parameters were calculated from the software to find the optimal fit to the data (producing the least variance). The extra sum of squares *F* test was used to determine if best fit values of selected parameters (PWC50 and *Hillslope*) differed significantly between SHADE and SOLAR datasets (Turner et al., 2015) i.e. is the error in the model reduced using specific parameters for each dataset vs using global/shared parameters. The alpha value for all significance testing was set as *p* < 0.05.

If a separate model was required for inclusion of the SOLAR data (based on a significant *F* test), correction factors were generated based on the difference in PWC% between the SHADE and SOLAR models. We then modelled the *difference* in PWC between the SHADE and SOLAR models to form correction factors based on the solar intensity. The correction factors were then used to estimate PWC during sunlight exposure, for heat indices that do not intrinsically account for radiation in their calculation (i.e. *T*_wb_, humidex, and heat index). The correction factor was formed based on the following template:

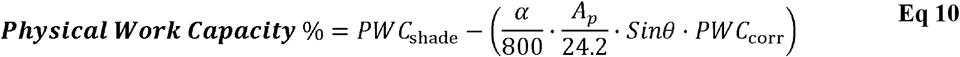

where *PWC*_shade_ calculates percent PWC in shaded conditions, *PWC*_corr_ is the correction factor observed in this experiment and 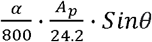 is a linear scaling factor which interpolates the SOLAR impact based on the present radiation. Equations for *PWC*_shade_ are available in our companion paper (Foster et al., 2021b); α is the SOLAR intensity in W·m^−2^. 800 W·m^−2^ was as the reference for the radiation level used in the present experiment. Using a SOLAR intensity of greater than 800 W·m^−2^ is possible but extrapolates beyond our empirical dataset rather than interpolating_;_ *A*_p_ is the projected area (the surface area of skin exposed to radiation), expressed as a percentage of total body surface area. 24.2% was used in the present study and represents a worst-case scenario (i.e., maximum possible surface area exposed to SOLAR). Sinθ is the sinus of the angle (θ in degrees) between the SOLAR beam and the surface onto which it projects. Sinθ values range from 0-1. Values for *A*_p_ can be calculated based on SOLAR altitude and azimuth (Underwood and Ward, 1966).

The predictive power of core and mean skin temperature for estimating PWC% was assessed using the same function as described in Eq 9, with the *x* value representing core or skin temperature. The relationship between these two thermometric variables and WBGT were assessed with a basic linear model.

## Results

### Physical work capacity

#### Dry climate

Significant main effects were found (Fig. 1 A,C) for condition (SOLAR-SHADED) and air temperature (*p* < 0.05), but not for clothing (*p* > 0.05). There was a significant 3-way interaction term, indicating that the impact of SOLAR on PWC varied with air temperature and clothing (*p* < 0.05). In low clothing coverage, there was no effect of SOLAR on PWC at 25°C (Δ=−1 ± 13%, ES = 0.09), or 35°C (−1 ± 9%, ES = 1.34). However, SOLAR decreased PWC at 40°C (−7 ± 5%, ES = 1.32), and 45°C (−37 ± 7%, ES = 5.23). In high clothing coverage, there was no significant effect of SOLAR on PWC at 25°C air temperature (−2 ± 13%, ES = 0.14). However, SOLAR reduced PWC at 35 (−6 ± 10%, ES = 0.57), 40 (−12 ± 8%, ES = 1.57), and 45°C (−15 ± 12%, ES = 1.29).

#### Humid climate

Main effects were found for condition and air temperature (*p* < 0.05), but not for clothing (*p* > 0.05) (Fig 1 B, D). There were no significant interaction terms, indicating that the impact of SOLAR on PWC was not dependent on the air temperature, or clothing (p > 0.05). In low clothing coverage, SOLAR decreased PWC at 25 (−12 ± 13%, ES = 0.95), 30 (−20 ± 15%, ES = 1.39), and 35°C air temperature (−15 ± 3%, ES = 4.96). In high clothing coverage, SOLAR reduced PWC at 25 (−9 ± 9%, ES = 1.03), 30 (−7 ± 7%, ES = 1.03), and 35°C air temperature (−9 ± 8%, ES = 1.11).

The individual level PWC responses in each condition are available in the supplementary material.

### Role of skin and core temperature in predicting physical work capacity

Figure 2 (panels A and B) shows the independent effect of skin and core temperature (average of each trial) on PWC. During physical activity at a fixed heart rate of 130 b·min^−1^ (which broadly reflects the self-pacing behaviour of workers), the skin, but not core temperature, is a predictor of PWC in the heat. While SOLAR results in a higher *T*_skin_, the predictive power of *T*_skin_ for estimating PWC does not appear to be impacted by addition of SOLAR as additional predictor in the equation, indicating that the same PWC relation holds true for both SOLAR and non-SOLAR conditions (R^2^ = 0.69). Figure 2 (panels C and D) shows the skin and core temperature response to increased environment heat stress, using WBGT as an example heat index. WBGT was a strong predictor of *T*_skin_, and the strength of the prediction was not impacted by the inclusion of SOLAR data (R^2^ = 0.82. Analysis demonstrated that for both models in Figure 2 (A and C), separate parameter values are not required for the shade and SOLAR datasets (*p* > 0.05). It is recommended that the equation for each model is taken from our recently published study using a larger dataset (Foster et al., 2021b). The primary aim of this analysis was to determine whether separate parameter values are required for the SOLAR data, which there were not.

**Figure 2.**
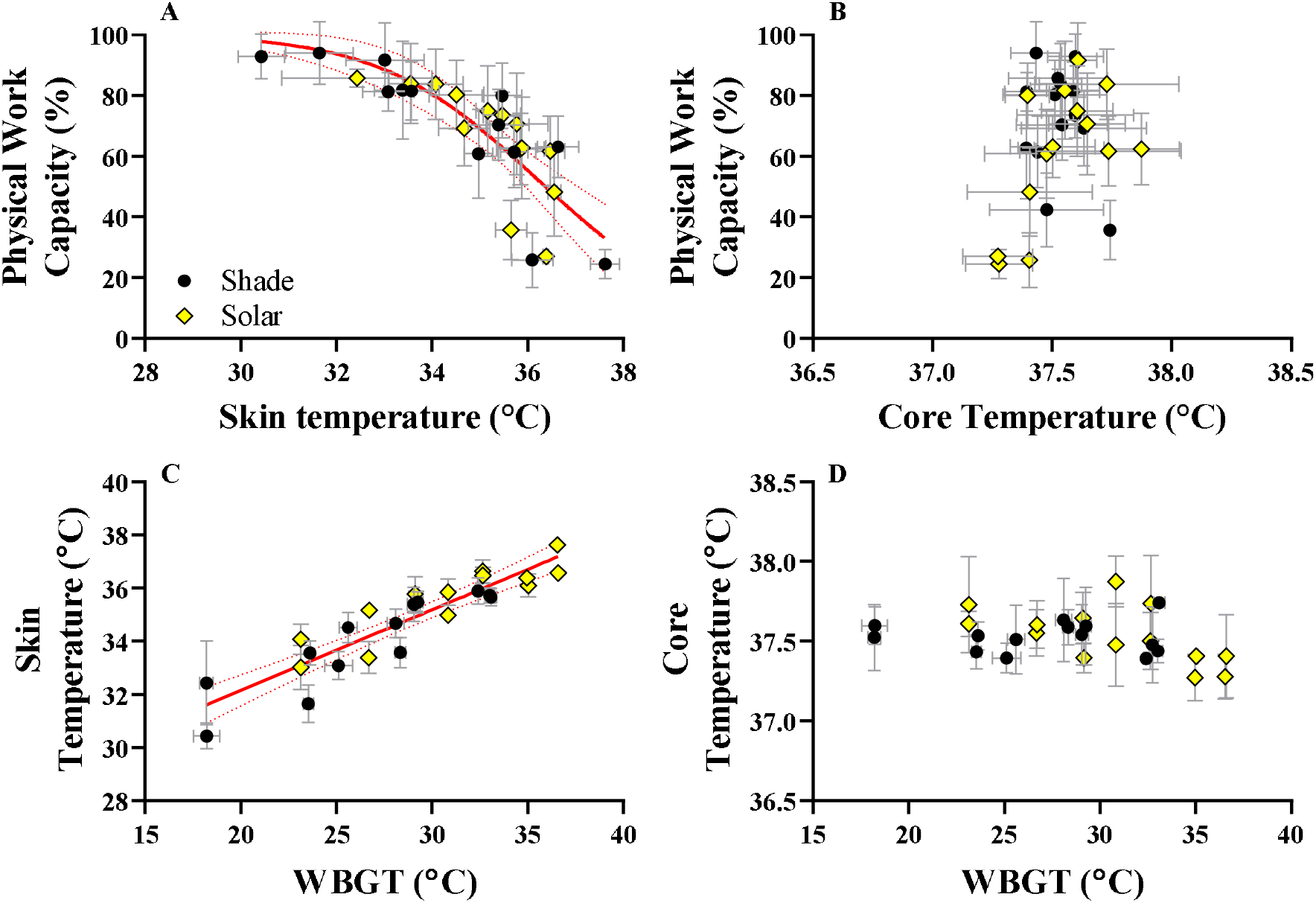
Increased skin, but not core temperature, is associated with PWC in the heat. Graphs A and B show the skin and core temperature (average over each trial) relationship with PWC, respectively. Graphs C and D show the skin and core temperature responses to varying WBGT while working at a fixed heart rate. Predictive power (R^2^) for models A and C were 0.68 and 0.82, respectively. Data is pooled for both clothing conditions. Black circles are SHADE trials, yellow diamonds are SOLAR trials. Dotted lines represent 95% confidence bands.

### Modelling physical work capacity with solar radiation

The WBGT and UTCI thermal indices account for SOLAR in their calculation, and therefore, a pooled model accounts well for both SHADE and SOLAR data. In both clothing conditions (Figure 3), changing the model parameters (PWC50 and Hillslope) for the SOLAR data did not improve the fit compared to a pooled model (*p* < 0.05). In other words, the WBGT and UTCI account well for the decrease in PWC with SOLAR, indicated by a rightward shift in their value under SOLAR exposure. This was true in both clothing conditions. Consequently, the inclusion of data with SOLAR does not negatively impact the error variance in the model, so there is no requirement to adapt the models already presented in our companion paper (Foster et al., 2021b). Since the *T*_wb_, Humidex, and Heat Index do not intrinsically account for thermal radiation, their values for a certain *T*_a_/Rh combination are unchanged with SOLAR, and the inclusion of SOLAR data decreases the predictive capacity of the model substantially. Hence, those heat stress indices require separate models for the SOLAR and SHADE (*p* < 0.05), since less residual variance was documented if separate models were produced for each dataset. This was true in both clothing conditions. For the heat stress indices that cannot use a pooled/global model, the *difference* between the SHADE and SOLAR functions were modelled from a gaussian expression. Those functions were used to form the correction factors (Table 2), where PWC predictions obtained from SHADE data are adjusted based on the solar load. These equations are based on the red area fill in Figure 3, which show the *difference* in PWC between SOLAR and SHADE conditions.

**Table 2.**
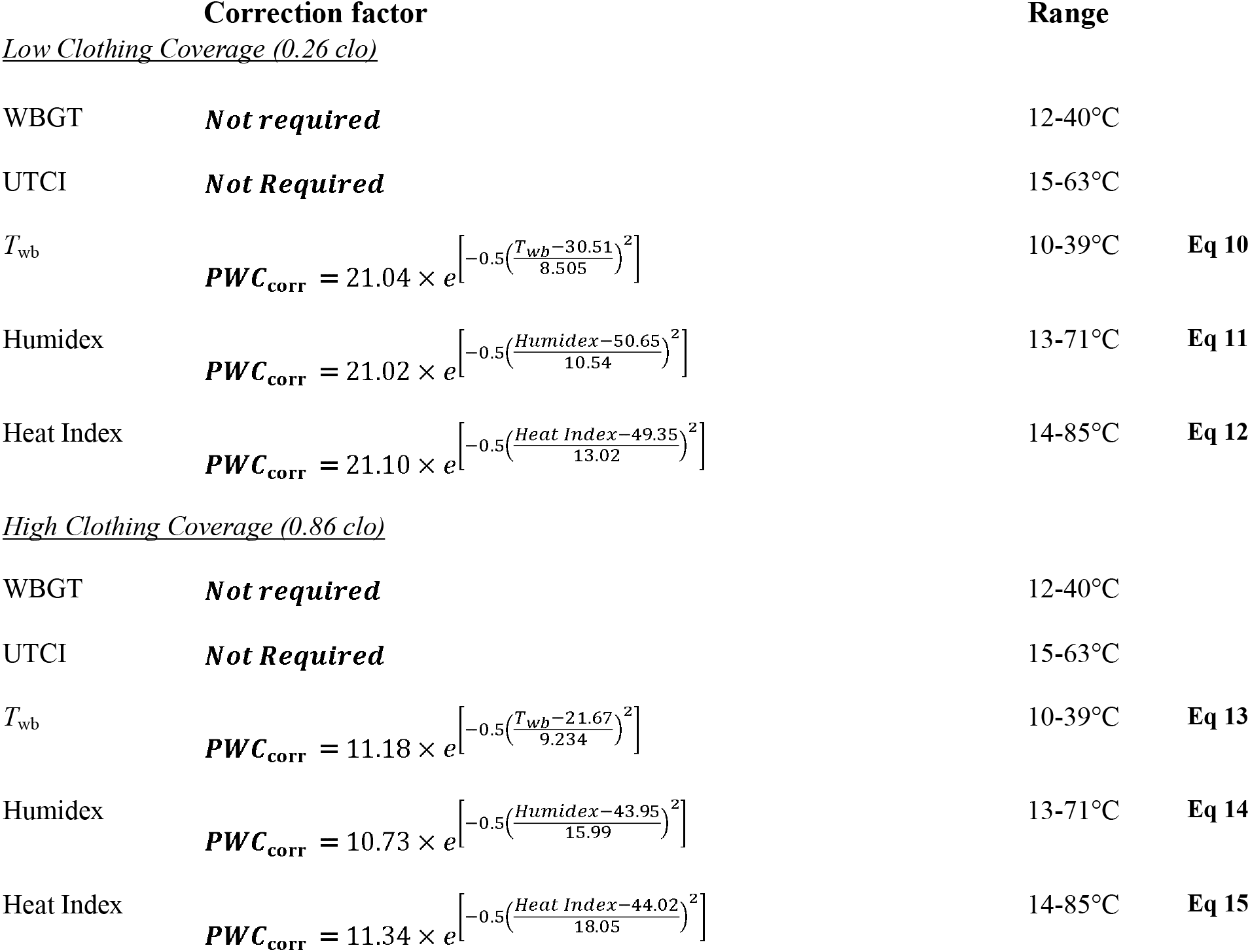
Correction factors for calculation of physical work capacity in outdoor working conditions with exposure to SOLAR. The values need to be subtracted from the shaded values in our companion paper as in Eq 10 (Foster et al., 2021b).

| **Correction factor** |  | **Range** |  |
| --- | --- | --- | --- |
| <u>Low Clothing Coverage (0.26 clo)</u> |  |  |  |
| WBGT | *Not required* | 12-40°C |  |
| UTCI | *Not Required* | 15-63°C |  |
| $T_{wb}$ | $PWC_{corr} = 21.04 \times e^{\left[-0.5\left(\frac{T_{wb}-30.51}{8.505}\right)^2\right]}$ | 10-39°C | **Eq 10** |
| Humidex | $PWC_{corr} = 21.02 \times e^{\left[-0.5\left(\frac{Humidex-50.65}{10.54}\right)^2\right]}$ | 13-71°C | **Eq 11** |
| Heat Index | $PWC_{corr} = 21.10 \times e^{\left[-0.5\left(\frac{Heat\ Index-49.35}{13.02}\right)^2\right]}$ | 14-85°C | **Eq 12** |
| <u>High Clothing Coverage (0.86 clo)</u> |  |  |  |
| WBGT | *Not required* | 12-40°C |  |
| UTCI | *Not Required* | 15-63°C |  |
| $T_{wb}$ | $PWC_{corr} = 11.18 \times e^{\left[-0.5\left(\frac{T_{wb}-21.67}{9.234}\right)^2\right]}$ | 10-39°C | **Eq 13** |
| Humidex | $PWC_{corr} = 10.73 \times e^{\left[-0.5\left(\frac{Humidex-43.95}{15.99}\right)^2\right]}$ | 13-71°C | **Eq 14** |
| Heat Index | $PWC_{corr} = 11.34 \times e^{\left[-0.5\left(\frac{Heat\ Index-44.02}{18.05}\right)^2\right]}$ | 14-85°C | **Eq 15** |

**Figure 3.**
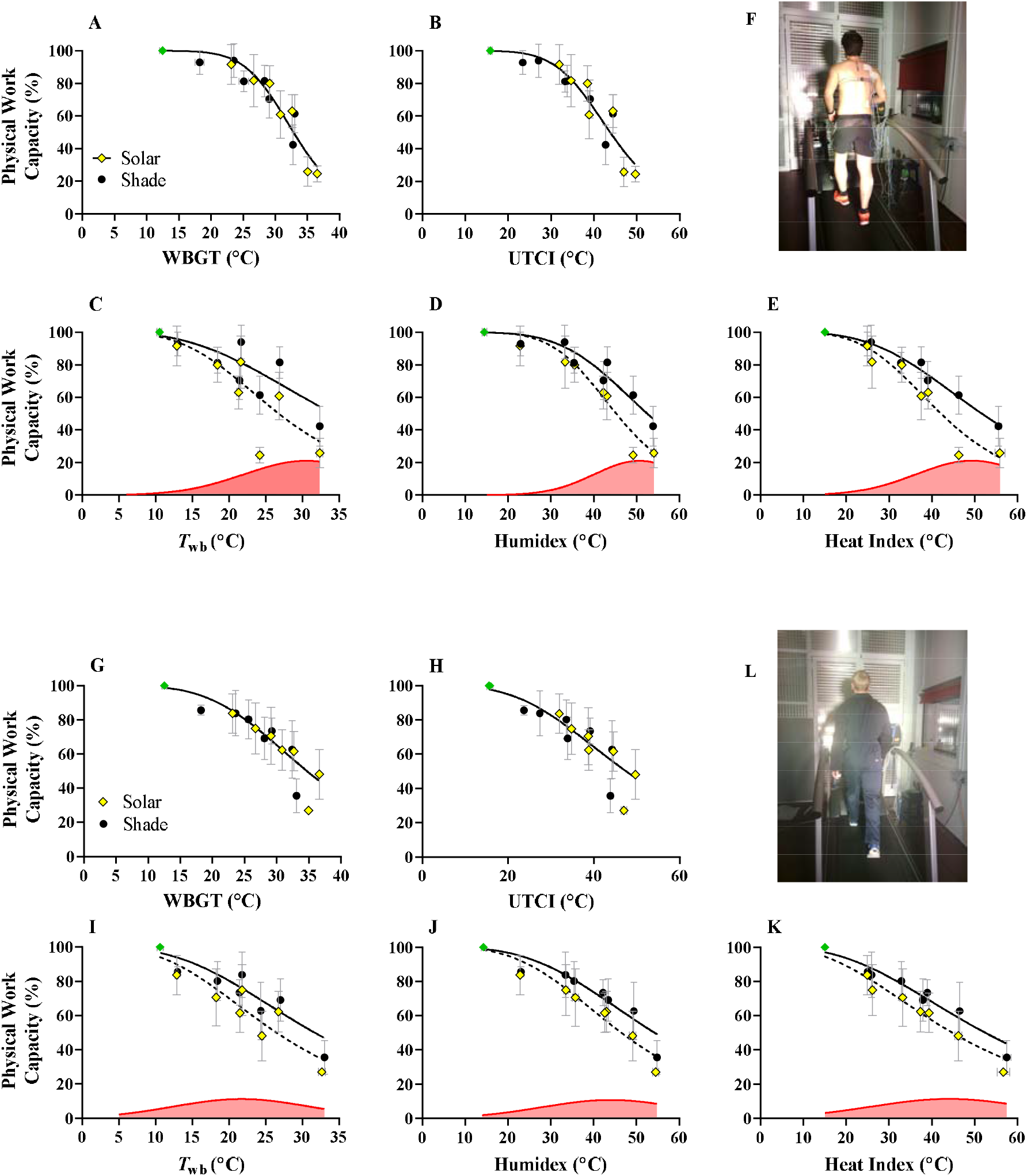
Physical work capacity (PWC) with low (A-F) and high clothing coverage (G-L). Black circles and yellow diamonds represent SHADE and SOLAR trials, respectively. For WBGT and UTCI, a pooled model is sufficient for predicting PWC even if solar data are included. For psychrometric wet-bulb (*T*_wb_), Humidex, and Heat Index, separate models are required if predicting PWC during SOLAR exposure. This was true for low and high clothing coverage. Here, the solid line and dotted line in figures represent SHADE and SOLAR data models, respectively. The *difference* in PWC (i.e. shade - solar) for models followed a guassian distribution, and is shown by the red area fill. These guassian models are shown in Table 4 and were required for the generation of correction factors, allowing computation of outdoor PWC for *T*_wb_, Humidex, and Heat Index. The clothing ensembles are shown in panels F and L.

These correction factors are for the full SOLAR load used in the experiment. Assuming a linear impact of the SOLAR intensity (based on the heat transfer into the body), for lower radiation levels, lower radiated areas and different SOLAR angles, the SOLAR impact can be scaled as:

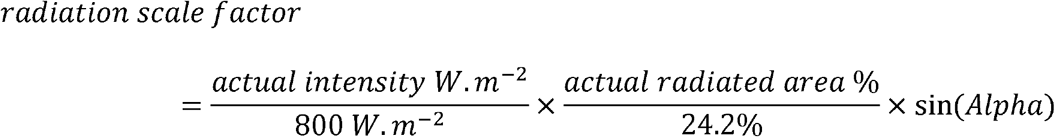

With Alpha=angle (in °) between radiation beam and projected surface; i.e. when radiation falls perpendicular onto surface, Alpha=90°.

### Example calculation

Below we provide an example calculation assuming a *T*_wb_ of 30°C, a SOLAR intensity of 600 W·m^−2^, projected over 15% BSA, and the radiation hitting the radiated surface at an angle of 45° (0.785 radians), with low clothing coverage.

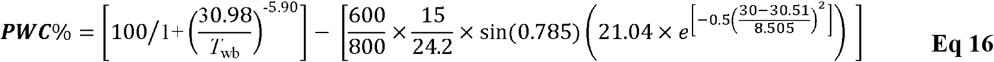

In this example, PWC is 48%. Without any SOLAR, PWC would be 55%. Note that the original/base model of PWC (the left term to the right of the equal sign) should be taken from our companion paper (Foster et al., 2021b).

## Discussion

This study builds on our earlier publication that determined Physical Work Capacity across a wide range of combinations of temperature and humidity (Foster et al., 2021b), but without SOLAR present. The primary aim of the present study was to determine the additional impact of SOLAR on human physical work capacity (PWC), with high and low clothing coverage and across a broad range of temperature and humidity combinations. The secondary aim was to evaluate whether PWC can be predicted outdoors with SOLAR using a variety of heat stress indices and finally whether our previously published functions linking PWC to the climate parameters can be used for both indoor (shade) and outdoor (SOLAR) work settings.

SOLAR reduced PWC by up to a further 20%, depending on the temperature, humidity, and clothing condition. We found the impact of SOLAR not to be ‘additive’ (Lloyd and Havenith, 2016), to the temperature, humidity, and clothing effects, but instead to show an interaction effect with these variables. With low clothing coverage in dry heat, SOLAR had a negligible impact on PWC when *T*_a_ ≤ 35°C, but PWC decreased exponentially due to SOLAR when *T*_a_ ≥ 40°C (Figure 1A). When high clothing coverage was adopted in the same climate types, SOLAR caused a consistent linear decrease in PWC when *T*_a_ ≥ 35°C (Figure 1C). In humid conditions, the impact of SOLAR was more consistent with an additive model in that the reduction in PWC caused by SOLAR was similar across all levels of *T*_a_ investigated, but slightly lower with high clothing coverage (Figure 1 B and D). Based on our data (comparing figs 2a to 2b), high clothing coverage only seems to protect against reductions in PWC caused by SOLAR when *T*_a_ ≥ 40°C, with the caveat that more protection from ultraviolet radiation may be required if low clothing coverage is adopted i.e., with sunscreen. The effect of sunscreen on human thermoregulatory function seems to also depend on the climate type (Connolly and Wilcox, 2000) and the composition of the ointment (Aburto-Corona and Aragón-Vargas, 2016), making any potential interaction difficult to predict at this stage.

In low clothing coverage (exposed skin), PWC was severely affected by SOLAR at 45°C/20% Rh, but the effect was substantially less with high clothing coverage for the *T*_a_ (Figure 1). Similar ‘protective’ effects of high clothing coverage during military marching in hot-dry conditions (43.3°C) with SOLAR have been described, evidenced by a substantial decrease in sweat output (Adolph, 1947). The response seems to be explained by the absolute skin temperature with SOLAR, which was greater at 45°C *T*_a_/20% Rh in low clothing coverage (37.6 ± 0.3°C) compared with high clothing coverage (36.6 ± 0.1°C). It is likely that the lower skin temperature in high clothing coverage is due to reflective properties of the clothing ensemble and the interruption of the direct radiation to the skin by absorption, which serve to reduce direct dry heat gain from SOLAR (Bröde et al., 2010a; Clark and Cena, 1978). However, it is worth noting that the beneficial impact of clothing depends on the colour and fabric properties (Bröde et al., 2010b; Nielsen, 1990). In dry heat when *T*_a_ ≤ 35°C, where no effect on PWC of adding SOLAR was observed, the added radiative heat gain was likely compensated for by increased sweat evaporation in low clothing coverage compared with high clothing coverage.

Mean skin temperature was a strong predictor of PWC in the heat, independent of clothing or exposure to SOLAR (Figure 2). In contrast, the core temperature had no predictive value for PWC at our chosen work rate. These findings are supported by prior work from our group (Foster et al., 2021b) and others (Ioannou et al., 2017; Jay et al., 2019), which implicates *T*_skin_ as the primary determinant of work capacity loss during occupational heat stress. Mechanistically, a study in mice showed that the strength of the afferent nervous system response is directly proportional to the absolute (not relative) skin temperature during heating (Ran et al., 2016), providing strong theoretical basis for our observations.

A fixed cardiovascular strain model was chosen as a proxy for self-paced physical work based on a plethora of field data in which workers can freely adjust their pace in hot climates (Bates and Schneider, 2008; Kalkowsky and Kampmann, 2006; Mairiaux and Malchaire, 1985; Miller et al., 2011; Wyndham, 1973). Discussing data from the South African gold mines (Wyndham, 1973), Vogt et al., (1983) observed that *“while productivity and oxygen consumption fell off with increasing wet-bulb temperature, heart rates remained constant around an average of 130-140 b*·*min^−1^.* A heart rate of 130 b·min was therefore chosen, which also represented an occupational intensity on the border of moderate to heavy physical work, as suggested by the World Health Organization (Andersen, 1978). Data from a follow up experiment with 6 one-hour work bouts on a single day, show that this heart rate represents a full day limit with participants feeling very fatigued upon cessation (Smallcombe et al., 2019). However, it is possible that e.g., for shorter periods than a day, a higher limit for the fixed heart rate is possible, and that with higher heart rates (i.e., higher workloads), core temperature and dehydration may become a more relevant predictor of the loss in PWC.

WBGT and UTCI are commonly used heat stress indices in biometeorology, and their values account for any change in mean radiant temperature (Havenith and Fiala, 2015). We show that, based on the change in mean radiant temperature with SOLAR (determined empirically by black globe temperature combined with *T*_a_ and Rh), the relative shift in the value of WBGT and UTCI predicts the reduction in PWC caused by SOLAR appropriately. Therefore, assuming that globe temperature is correctly measured (spatially and allowed to equilibrate), the WBGT and UTCI can accurately predict PWC in both shaded and unshaded conditions with a single equation. The validity of models that use WBGT to predict PWC (Dunne et al., 2013; Kjellstrom et al., 2018) is therefore not reduced if also applied to outdoor work settings. In contrast, the *T*_wb_, Humidex, and Heat Index do not intrinsically account for any change in mean radiant temperature, highlighting the importance of context if such indices are linked with human physiology or survival. Correction factors for PWC are provided if such indices are to be used for outdoor work with SOLAR, as shown in Table 2. The equations reported in this study have immediate applicability for those studying the impact of hot weather on PWC, especially in outdoor settings.

## Limitations

There are several limitations of the present study that should be considered. Firstly, it is unclear if the general conclusions about SOLAR are accurate during exposure to a full working day, in contrast to the 1-hour work bout used in our study. Preliminary data from our lab (Smallcombe et al., 2019) indicates minimal impact of work duration on PWC until WBGT reaches 36°C, which is only surpassed at 45°C/20% Rh and 35°C/80% Rh with the addition of SOLAR. Finally, a constant intensity of 800 W·m^−2^ SOLAR was chosen for this study. Ideally, different intensities would be measured to improve impact analysis in different regions or times of day. The choice of radiation intensity and radiated surface area represented a worst-case scenario, simulating outdoor work under direct sunlight at the hottest part of the day (Monteith and Unsworth, 1990). The correction factors are thus based on the assumption that the radiation level can be described as a linear impact between our shade and 800 W·m^−2^ condition. While from a heat balance modelling perspective, this is deemed plausible, research should verify the validity of this assumption.

## Conclusions

Addition of solar radiation to climatic heat stress reduced PWC by up to a further 20%, above the SHADE condition depending on the temperature, humidity, and clothing condition. We observed an interactive effect of solar radiation on physical work capacity, depending on clothing coverage and the climate. WBGT and UTCI account well for the impact of solar radiation on physical work capacity. Correction factors are available if using *T*_wb_, humidex, or heat index for the prediction of physical work capacity, in solar radiation conditions.

